# Nonnegligible role of warming-induced soil drying in regulating warming effect on soil respiration

**DOI:** 10.1101/273573

**Authors:** Enzai Du

**Affiliations:** State Key Laboratory of Earth Surface Processes and Resource Ecology, Faculty of Geographical Science, Beijing Normal University, Beijing, 100875, China

## Abstract

Based on results of a 26-year soil warming experiment (soil temperature being elevated by 5 °C) in a Harvard hardwood forest, Melillo et al. demonstrated a four-phase pattern of long-term warming effect on soil respiration, while the mechanisms were not fully elucidated because they neglected the indirect effect due to warming-induced soil drying. By showing a significant correlation between precipitation anomaly and inter-annual variation of warming effect on soil respiration, we suggest a nonnegligible role of warming-induced soil drying in regulating the long-term warming effect on soil respiration. Our analysis recommends further efforts to consider both the direct and indirect (i.e., warming-induced soil drying) warming effects to gain more in-depth understanding of the long-term soil C dynamics.

---

Understanding the long-term effect of climate warming on soil respiration is a prerequisite for the projection of future change in global soil carbon (C) pools and their consequent feedbacks to climate change. Based on results of a 26-year soil warming experiment (soil temperature being elevated by 5 °C) in a Harvard hardwood forest, Melillo et al. (1) proposed a hypothesis of four-phase pattern of soil warming effect on soil respiration (ΔRs = Rs_Heated_-Rs_Control_) and attributed it to soil substrate and microbial changes. However, the mechanisms were not fully elucidated because the authors improperly neglected the indirect effect due to warming-induced decline in soil moisture (Fig. 1).

**Figure 1.**
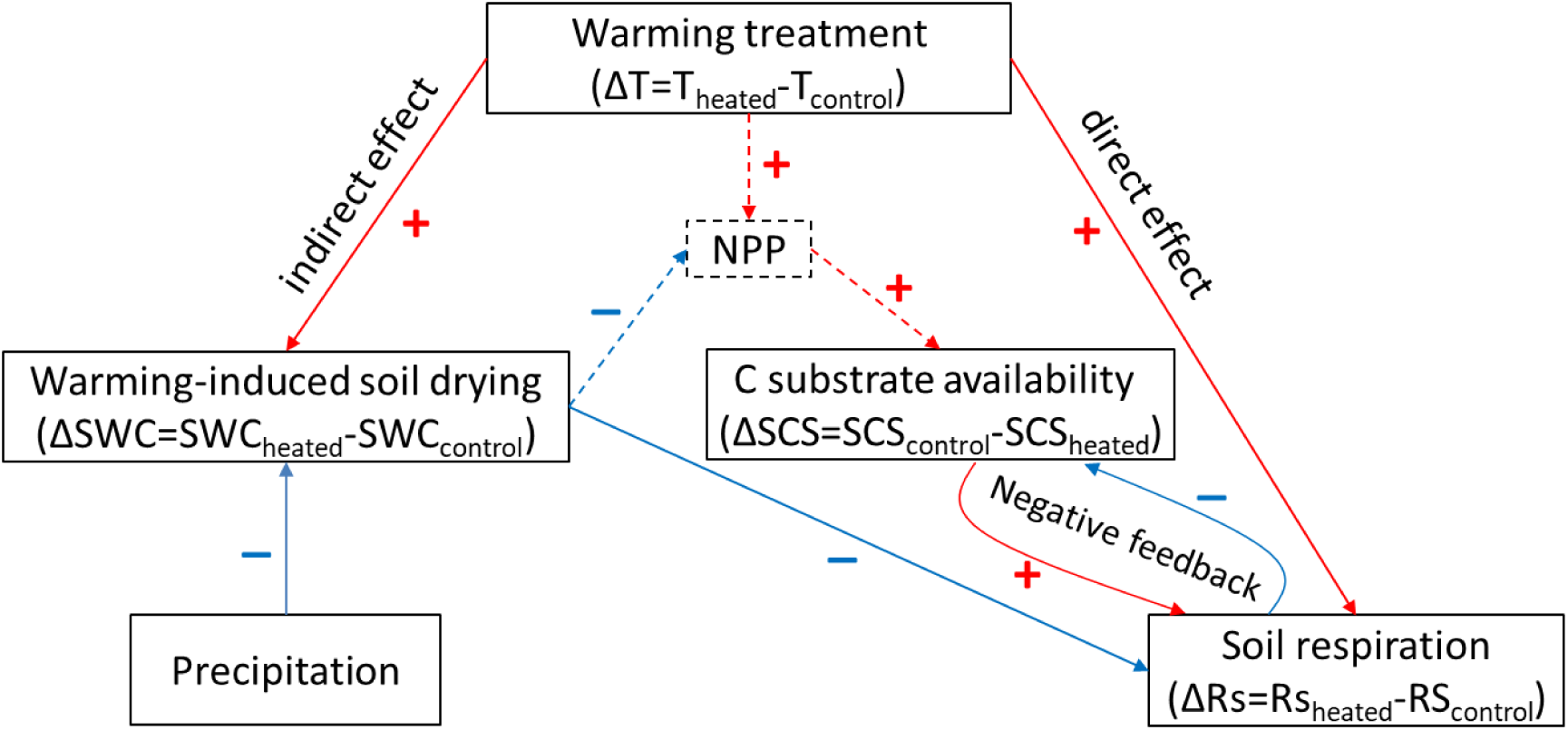
A conceptual framework showing the direct and indirect warming effects on soil respiration. Precipitation buffers warming-induced soil drying and thus has an positive effect on inter-annual variation of ΔRs. Negative feedbacks exist between C substrate availability and respirational C loss. Current experiment failed to track the direct and indirect warming effects on forest primary productivity (NPP) and consequent fresh soil C inputs due to a small plot size (6 x 6 m^2^) (as indicated by dashed lines). The symbols “+” in red and “–” in blue indicate positive and negative effects, respectively.

Experimental results have evidenced an essential role of soil moisture in regulating the warming effect on soil respiration (2-5). Lower soil moisture can reduce ΔRs, and even leads to negative ΔRs, when soil moisture becomes extremely limiting to plant roots and soil microbial organisms (Fig. 1; 3-6). In addition to the direct warming effect (1), warming-induced soil drying (2-6) may indirectly reduce ΔRs, especially in drier years (Fig. 1; 2-5). Moreover, soil moisture stress has been evidenced to reduce temperature sensitivity of soil respiration (7, 8). Warming-induced soil drying may have also contributed to a decrease in temperature sensitivity of soil respiration in heated plots, being observed to occur consistently across all four phases during the long-term soil warming experiment in Harvard forest [see Fig. 3 in Melillo et al. (1)]. Unfortunately, we cannot re-analyze the role of warming-induced soil drying due to unavailable soil moisture data (1).

As a primary source of soil moisture, precipitation is expected to buffer the soil moisture stress and result in a positive effect on inter-annual variation of ΔRs (Fig. 1). By retrieving data on total annual precipitation (TAP) anomaly (showing no significant temporal trend, *p=0.94*) and ΔRs from Melillo et al. (1), we tested the effect of TAP anomaly on inter-annual variation of ΔRs. As ΔRs showed a significant decrease over time (ΔRs = –6.1547*Year + 12402, adjusted *R*^*2*^ = *0. 29, p=0.005*) due to long-term microbial C substrate depletion (Fig.1; 1, 9, 10), we thus used detrended (i.e., residual) ΔRs [being calculated as, measured ΔRsi – (–6.1547*Yeari +12402), where i indicates a specific year] to indicate the inter-annual variation of ΔRs. As expected, we found a significantly positive effect of precipitation anomaly on the inter-annual variation of ΔRs (slope = 0.15 g C m^−2^ year^−1^ per mm; adjusted *R*^*2*^ = *0.17, p=0.03*; Fig. 2). Our result suggests a significant role of soil moisture in regulating the warming effect on soil respiration. It further implies that warming-induced soil drying stress has very likely exerted a negative effect on the response of soil respiration to long-term warming treatment in Harvard forest.

**Figure 2.**
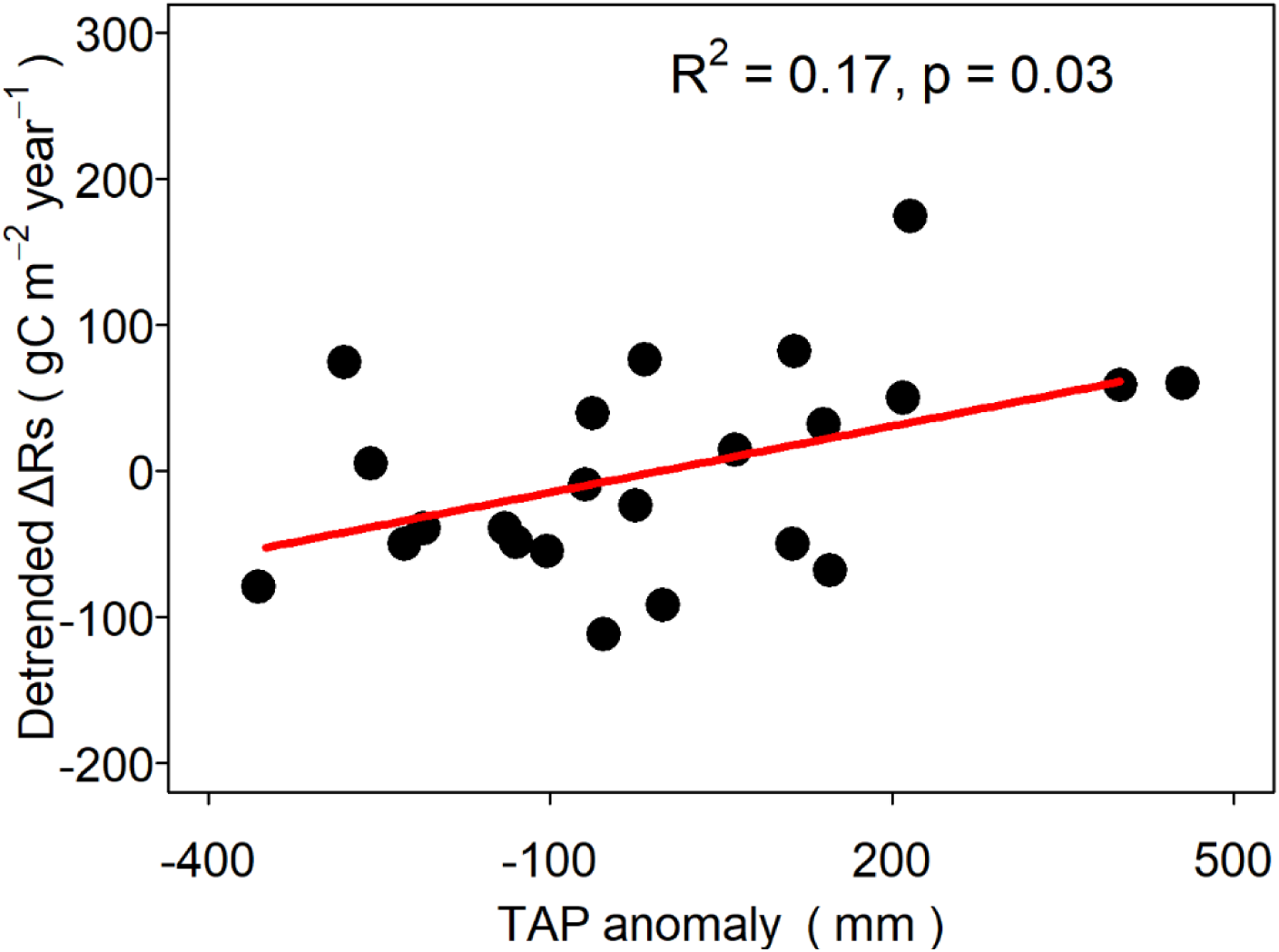
Significant effect of total annual precipitation (TAP) anomaly (mm) on the inter-annual variation of warming-induced ΔRs (g C m^−2^ year^−1^). Because ΔRs showed a significant decrease over time (ΔRs = –6.1547*Year + 12402, adjusted R^2^= 0.29, p=0.005) due to soil C substrate depletion, we used detrended residual [being calculated as, measured ΔRsi–(–6.1547*Yeari + 12402), where i indicates a specific year] to indicate the inter-annual variation of ΔRs. Note that our analysis excluded data for the years 1995, 2005, 2010 due to warming system failure.

The long-term experiment, with each plot covering a small area of 6 × 6 m^2^ (1), generally represented a scenario of soil C depletion under soil warming, while it failed to demonstrate the story in a whole C cycle perspective. The small plot size was not able to track the direct and indirect (i.e., warming-induced soil drying) warming effects on forest primary productivity (Fig. 1), which could consequently provide fresh C substrates for soil respiration (11). In that case, negative feedbacks between C substrate limitation and respirational C loss likely dominate a long-term reduction of ΔRs (9, 10). By inducing a reduction of soil moisture (2-6, 12, 13), soil warming also indirectly regulates ΔRs via inducing changes in the composition and activity of microbial community (5, 14). For instance, warming-induced soil moisture stress may have contributed substantially to the declining trends of ΔRs during the relatively drier periods 1997-2002 and 2015-2016 [see Fig. S2 in Melillo et al. (1)].

As the indirect effect due to warming-induced soil drying stress has not been addressed properly, the mechanisms of the four-stage pattern of long-term warming effect on soil respiration are likely not fully elucidated (1). Moreover, Melillo et al. (1) assumed that microbial respiration accounted for a constant proportion (approximately 2/3) of soil respiration, while they ignored temporal changes in the microbial proportion over the four-phase pattern of soil respiration. Therefore, in-depth understanding of the long-term warming effect on soil C dynamics calls for further efforts to consider both the direct and indirect (i.e., warming-induced soil drying) effects in a whole C cycle perspective at an ecosystem scale.

## Acknowledgements

This work was supported by the National Natural Science Foundation of China (Nos. 41630750 & 31400381) and State Key Laboratory of Earth Surface and Resource Ecology (No. 2017-ZY-07).

